# Two *Pellioditis* biocontrol nematode species infect *Ariolimax columbianus*, the Pacific banana slug, and increase mortality in laboratory infectivity trials

**DOI:** 10.64898/2026.05.14.725190

**Authors:** Emily R. Taylor, Indira Kulkarni, Dana K. Howe, Casey H. Richart, Rory J. Mc Donnell, Dee Denver

**Affiliations:** Department of Integrative Biology, Oregon State University, Corvallis, OR 97331, USA; Department of Crop and Soil Science, Oregon State University, Corvallis, OR 97331, USA; Department of Forest Ecosystems & Society, Oregon State University, Corvallis, OR 97331, USA

## Abstract

Gastropods are a highly diverse and often overlooked taxonomic group of significant ecological and economic importance. Some terrestrial gastropods are critical pests of commercial agriculture and home gardens worldwide. Malacopathogenic nematodes offer an effective biological control method of managing pest slugs and snails as a ‘natural enemy’. *Pellioditis* (syn. *Phasmarhabditis*) *hermaphrodita* and *Pellioditis* (syn. *Phasmarhabditis*) *californica* are two species of biocontrol nematodes that have been commercialized, sold as Nemaslug® and Nemaslug® 2.0 respectively on three continents. Although there is interest in bringing Nemaslug® products to the US, they are currently not permitted due to limited knowledge on their North American distribution and effects on non-target and native species. In this study, we investigated the impact of *P. hermaphrodita* and *P. californica* on *Ariolimax columbianus* across two slug-host life stages, in laboratory infectivity assays. The objectives were to 1. determine whether *P. hermaphrodita* and *P. californica* nematodes impact survival of *A. columbianus*, and 2. evaluate whether there are differential effects on survival in juvenile and adult life stages of *A. columbianus*, in laboratory infectivity trials. We found that *P. hermaphrodita* caused significant mortality in *A. columbianus* with 100% mortality observed in both juvenile and adult slug hosts. The *P. californica* treatment had significant effects on the juvenile *A. columbianus* group only, with 80% mortality. By contrast, only 16% of unexposed control juveniles and 4% of control adult slugs died during the experiment. These results indicate that *P. hermaphrodita* and *P. californica* are lethal to the native, non-target Pacific banana slug (*A. columbianus*) under laboratory conditions, with mortality differing between juvenile and adult host life stages. Given the ecological importance of *A. columbianus*, these findings raise concerns for potential non-target effects of *P. californica* and *P. hermaphrodita* on terrestrial gastropod communities and emphasize the need for testing biocontrol agents against multiple life stages.

## Introduction

Maintaining biodiversity is crucial to an ecosystem’s overall health and stability [1]. Historically, in the world of nonmarine mollusk research, scientific focus has often centered on the eradication of pest species, such as the grey field slug, *Deroceras reticulatum*, an invasive estimated to cause $60 million in losses to the state of Oregon’s grass seed industry annually [2]. Consequently, less attention is paid to the many important native species that make up a large part of natural ecosystems [3]. Many species of native terrestrial gastropods (land slugs and snails) play an important role in the ecosystem as detritivores, aiding in nutrient cycling and decomposition which benefits soil health [4]. Insufficient knowledge of the ecology of many native gastropod species hinders efforts to effectively conserve their biodiversity, with mollusks comprising roughly one-third of all recorded animal species extinctions since 1500 CE [5].

In addition to a lack of knowledge, there is a concerning legacy of human-implemented biological control (‘biocontrol’) strategies involving terrestrial gastropods that have resulted in ecological disaster for native species [3, 6-8]. A well-known example is the rosy wolfsnail (*Euglandina rosea*), a predatory terrestrial snail introduced to Hawaii in the 1950s to control pest species. Rather than controlling target pest populations, *E. rosea* became established in the local ecosystem and has now been widely implicated in the extinction of multiple native Hawaiian snail species, as well as additional extinctions across the Pacific Islands [6-8]. Given this unfortunate history, thorough host range testing of native gastropod species has been identified as a crucial step in assessing effectiveness and safety of biocontrol products [6].

Malacopathogenic nematodes, such as *P. hermaphrodita* and *P. californica*, offer a modern biocontrol solution for eliminating pest slugs from crops and gardens. The nematode *P. hermaphrodita* is a commercialized biocontrol species sold as Nemaslug® in Canada, Europe, and Kenya [9], with the purpose of eliminating target pests such as *D. reticulatum*. The nematode *P. californica* is also a gastropod-killing species that has shown increasing potential as a replacement for *P. hermaphrodita* and is currently sold as the new and improved Nemaslug® 2.0 [9, 11]. Both nematode species were utilized in this study.

While not currently permitted for sale in the US, there is increasing interest in bringing Nemaslug® to the US market as a natural alternative to current chemical control methods such as metaldehyde pellets, which can be ineffective and potentially harmful to the environment [10, 12]. The lethality of *P. hermaphrodita* to pest *D. reticulatum* is well-studied [9], in addition to potential non-target species in Europe [13], but there is a lesser volume of research on non-target species native to the USA. Recently, *P. hermaphrodita* has been shown to be lethal to the Pacific Northwest native snail *Monadenia fidelis*, the Pacific sideband, offering one example of a non-target effect in a US ecosystem [14]. As identified by Christensen et al. [6], expanded host range testing of non-target species is critical in understanding the impact of biocontrol strategies on the ecosystem.

Previous infectivity studies, primarily focused on species of economic relevance, have found *P. hermaphrodita* to be lethal to a diverse range of terrestrial gastropods, including slugs belonging to the families Arionidae, Limacidae, and Milacidae [9, 11, 16-19], as well as several snail species [9, 13, 19-21]. As mentioned previously, recent work has also found a native non-target in the US, *M. fidelis*, to be susceptible to *P. hermaphrodita* [14]. *P. californica* has been found to cause mortality in the slug *D. reticulatum* and the White garden snail (*Theba pisana*), the amber snails (*Succinea* spp.), and neonate garden snails (*Cornu aspersum*) [11, 20-23]. In some species, the effectiveness of nematode infection varies depending on the life stage of the host organism, such as in the Portugese slug (*Arion lusitanicus*) where individuals are susceptible to *P. hermaphrodita* at the juvenile stage, but adults are unaffected [17, 24]. To date, no slugs within Ariolimacidae have been tested.

The non-target focus of this study is the Pacific banana slug (*Ariolimax columbianus*) (Fig 1). *A. columbianus* is a North American Pacific coast native gastropod that can be found ranging from Alaska to northwestern California. They are the largest gastropod species found on the North American continent [15]. In addition to their vital role in the ecosystem as detritivores and decomposers, banana slugs are recognized as familiar cultural icons of the North American West coast. Their status as a native, non-target, and well-known species makes them a suitable candidate for non-target testing in this study.

**Figure 1.**
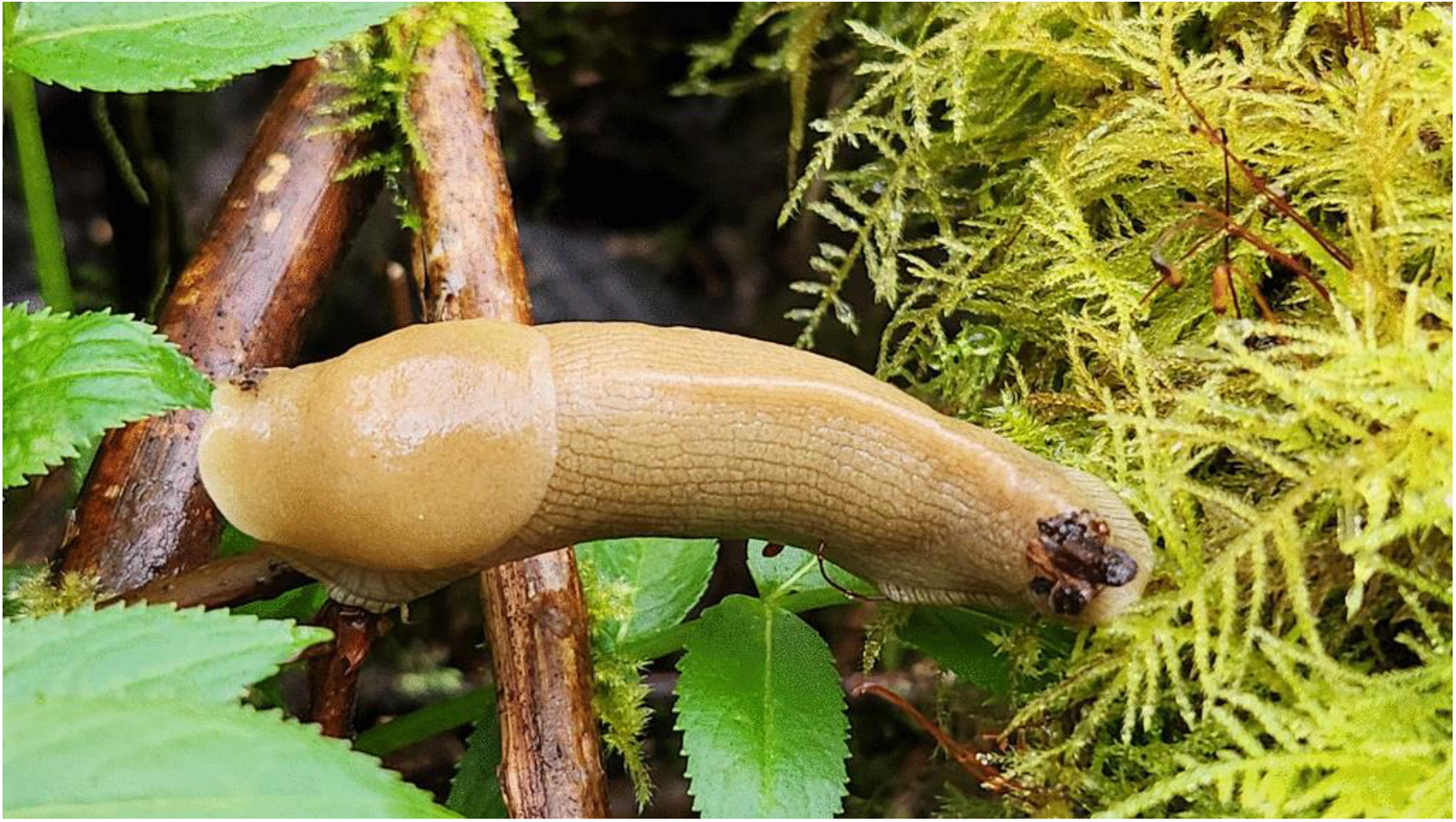
*A. columbianus* in its natural ecology on the Oregon coast. Photo by C.H. Richart.

This study investigated the impacts of *P. hermaphrodita* and *P. californica* on the non-target slug species *A. columbianu*s, in laboratory infectivity trials. Both juvenile and adult host life stages were tested for susceptibility to nematode infection. Mortality rates between life stages were expected to differ, as shown by Grimm [24], where the smallest weight class of the pest slug *A. lusitanicus* showed the highest nematode-induced mortality. This pattern is consistent with the observation that *M. fidelis* with larger-diameter shells showed higher survival rates in Denver et al. [14]. The difference in survival between life stages has not yet been tested in a non-target US-native slug species and provides key insight into how *P. hermaphrodita* and *P. californica* might affect non-targets. The results of this experiment inform our understanding of the potential impacts of biocontrol nematodes on non-target slugs in the Pacific Northwest, an area with rich terrestrial gastropod diversity.

## Materials and Methods

### Slugs and Nematodes

*Ariolimax columbianus* samples were collected along the Oregon coast from March 2024 to April 2024. As per Oregon Department of Agriculture policies, permits were not required to collect *A. columbianus* (a non-threatened terrestrial gastropod species) from public lands. Individuals were identified as juveniles or adults based on weight, with juveniles ranging from 0.22-2.67g, and adults ranging from 6.60-30.7g (Fig 2). These ranges were comparable to weight group classifications described in Pearson et al. [25]. Weight range outliers were not included in this study. All specimens were kept in plastic containers and fed organic sweet potatoes, as outlined in Mc Donnell et al. [11], before and during infectivity trials. Slug specimens were brought into the lab and kept in an incubator at 18℃ with a 12-hour photoperiod until trials began.

**Figure 2.**
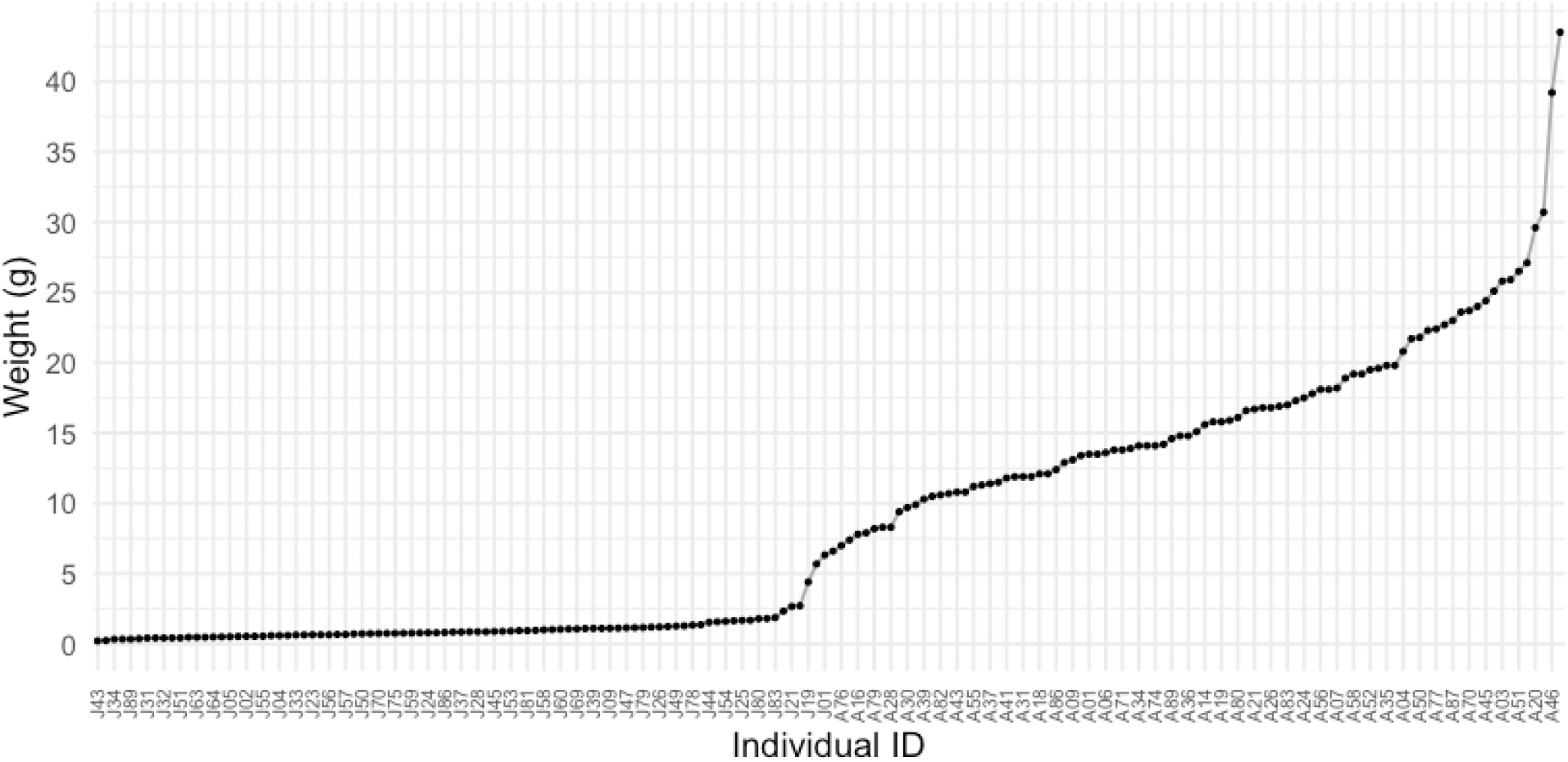
*Ariolimax columbianus* rank-ordered weight distribution. Weight in grams (y-axis) of each *A. columbianus* individual (x-axis) collected for the trial (n=178), arranged in ascending order to illustrate size variation. Individual IDs (x-axis) indicate juvenile or adult classification followed by a unique numerical identifier (‘J’ for juveniles; ‘A’ for adults).

The Oregon-derived *P. hermaphrodita* strain DL309 and the Washington-derived *P. californica* strain DL320 were used in this study. We also note here that while *P. hermaphrodita* and *P. californica* were previously classified in the *Phasmarhabditis* genus, recent work has led to its dissolution, with ‘*Pellioditis*’ now replacing ‘*Phasmarhabditis’* as the genus for this group of nematode species [26]. These nematodes were grown, collected, and washed using the modified White traps protocol to select for infective juvenile stage nematodes [27]. Infective juvenile stage nematodes (IJs) of each strain were aliquoted into individual 5mL suspensions at the BASF recommended application rate of 30 IJs/*cm*2.

### Infectivity Assays

Infectivity assays took place in plastic containers with a bottom layer of autoclaved soil, in two different container sizes corresponding to life stage. Juvenile *A. columbianus* were housed in round plastic containers 64 *cm*2 in area with 25g of autoclaved soil, as described in Denver et al. [14]. Adult *A. columbianus* were housed in rectangular plastic containers 435 *cm*2 in area with 170g of autoclaved soil. This trial included three experimental groups: *P. hermaphrodita* (strain DL309) treatment, *P. californica* (strain DL320) treatment, and negative control treatment where no nematodes were introduced. Each treatment was applied to one group of adult slugs and one group of juvenile slugs. Nematodes were suspended in 5 mLs of DI water and applied evenly across the soil surface in each treatment container at the BASF recommended application rate of 30 IJs/*cm*2. Control group containers received 5mL of DI water without nematodes. Five slugs of the same life stage were added to each container, and a total of five replicates (five containers) per treatment group were used, for a total of 30 assay arenas. Slugs were fed organic sweet potato every two days, with uneaten food removed after 24 hours.

Infectivity trial containers were maintained in a Thermo Precision Model 818 growth chamber at 18℃ with a 12-hour photoperiod. During the infectivity trial, containers were checked daily and monitored for mortality until day 38 of the experiment, in which the trial was ended. Mortality was assessed following a modified version of the procedure outlined in McDonnell et al. [11], including assessment of physical cues such as discoloration, body position (e.g. laying on their side), and lack of behavioral response to eyestalks stimulated with a metal spatula.

### Statistical Analyses

Slug survival data was initially visualized by plotting total percent mortality over time (38 days total) for each treatment group. To formally assess survivorship differences, Kaplan-Meier survival curves were generated using the survival package in RStudio (version 2025.05.1+513) for each treatment group. This non-parametric method of analysis allowed for right-censored observations, i.e. slugs that remained alive by the end of the trial, to be accounted for. Pairwise differences in survivorship between treatment groups were evaluated using log-rank tests, with an alpha level of 0.05 and p-values adjusted for multiple comparisons using the Bonferroni correction (see Table 1).

**Table 1.** Adjusted pairwise log-rank test p-values for survival comparisons between treatment groups.

| Treatment | Adult Ctrl | Adult Ph | Adult Pc | Juv Ctrl | Juv Ph | Juv Pc |
| --- | --- | --- | --- | --- | --- | --- |
| Adult Ctrl |  |  |  |  |  |  |
| Adult Ph | 2.7e-11* |  |  |  |  |  |
| Adult Pc | 1.0 | 6.6e-12* |  |  |  |  |
| Juv Ctrl | 1.0 | 2.7e-11 | 1.0 |  |  |  |
| Juv Ph | 2.6e-11 | 5.8e-13* | 5.8e-13 | 8.2e-13* |  |  |
| Juv Pc | 5.4e-07 | 1.0 | 6.8e-07* | 4.6e-06* | 0.16 |  |

Tests were adjusted for multiple comparisons using the Bonferroni correction. Significant differences (p<0.05) are indicated with an asterisk.

## Results

The results of the laboratory infectivity assay demonstrated that both *Pellioditis* nematode species significantly increased mortality in at least one life stage of *A. columbianus*, as compared to negative controls (Table 1). During the trial, nematodes could be seen actively infecting *A. columbianus* specimens, observed through a dissecting microscope on the outer surface of the mantle and eventually in large numbers on the slug cadaver (Fig 3). *P. hermaphrodita* was found to have significant effects on survival in both life stage groups (juvenile and adult), while *P. californica* only significantly impacted survival in the juvenile group. Overall, *P. hermaphrodita* killed host slugs at a faster rate than *P. californica* (Fig 4, 5). The first slug mortalities occurred on Day 5, with one slug death each in the juvenile *P. hermaphrodita* group and the juvenile *P. californica* group (Fig 5). The juvenile *P. hermaphrodita* treatment was the first to experience complete mortality in 16 days. The adult *P. hermaphrodita* group also reached complete mortality, at 27 days. Although death occurred in both, neither the juvenile nor adult *P. californica* treatment groups reached complete mortality by the conclusion of the trial.

**Figure 3.**
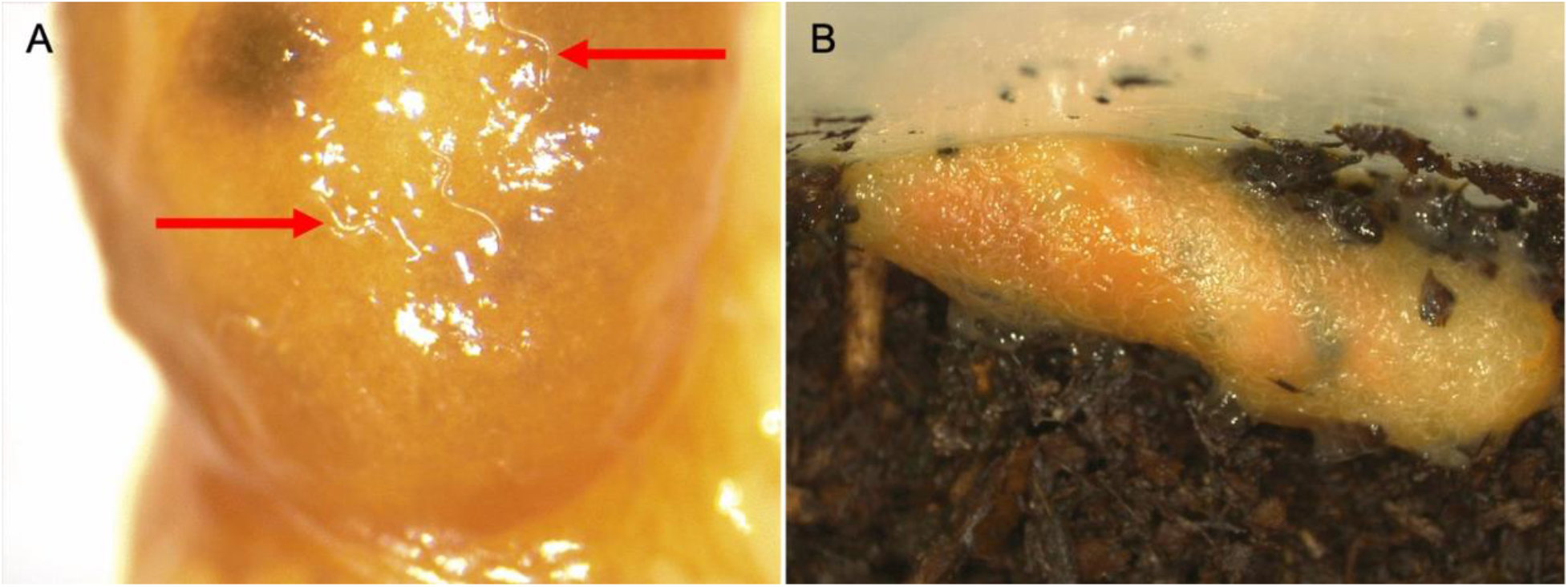
*P. hermaphrodita*-infested juvenile *A. columbianus* individual from the laboratory infectivity trial. **(A)**. Several *P. hermaphrodita* nematodes are visible on the outer surface of a juvenile *A. columbianus* mantle under a dissecting microscope, as indicated by red arrows. **(B)**. Thousands of *P. hermaphrodita* nematodes seen on the cadaver of a juvenile *A. columbianus*.

**Figure 4.**
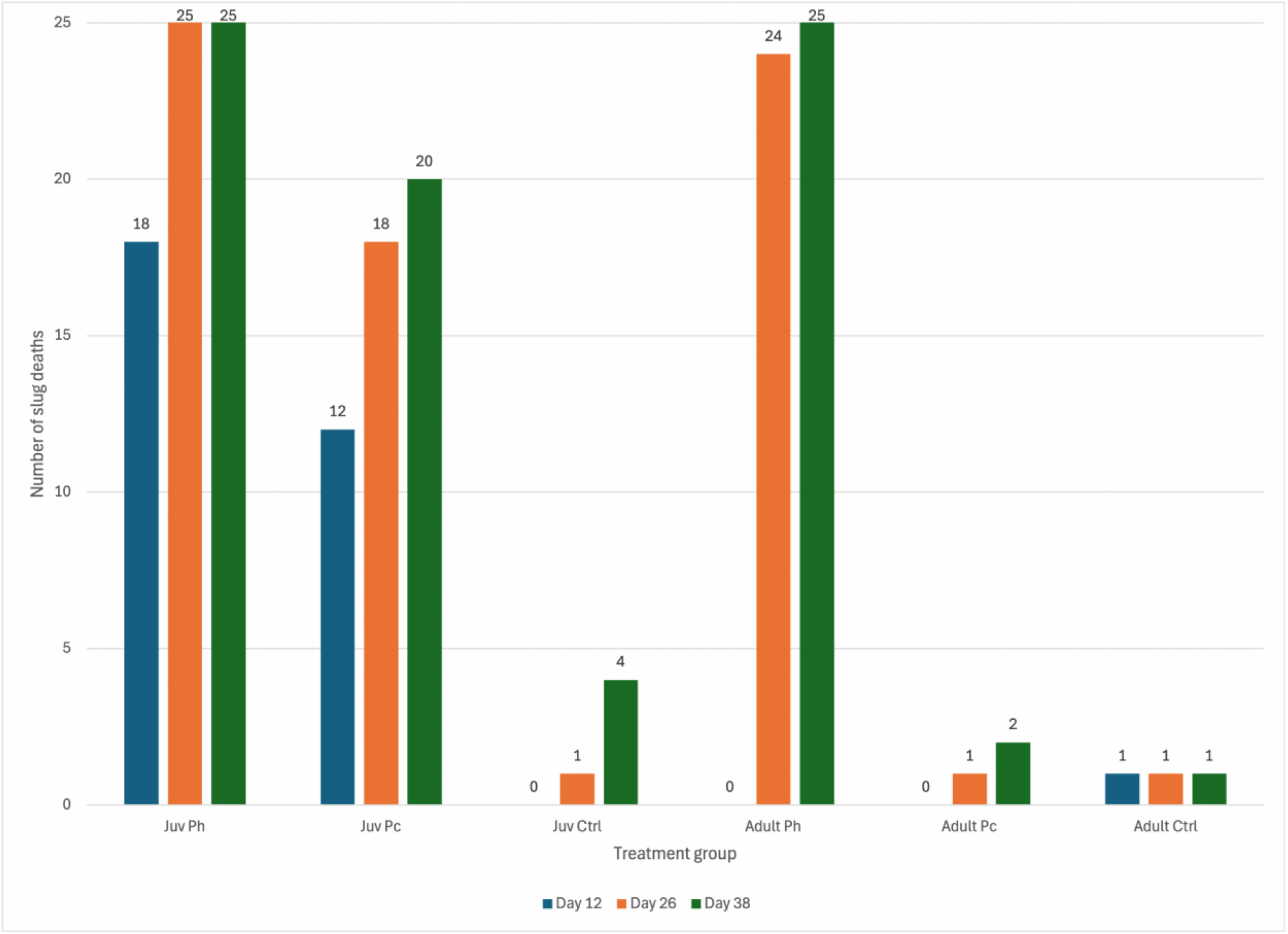
*A. columbianus* mortality at three time points during the infectivity trial (Day 12, Day 26, and Day 38). Raw count numbers depict the number of dead slugs per treatment group across three days of the trial (Day 12, blue; Day 26, orange; Day 38, green), out of a total of 25 slugs per group.

**Figure 5.**
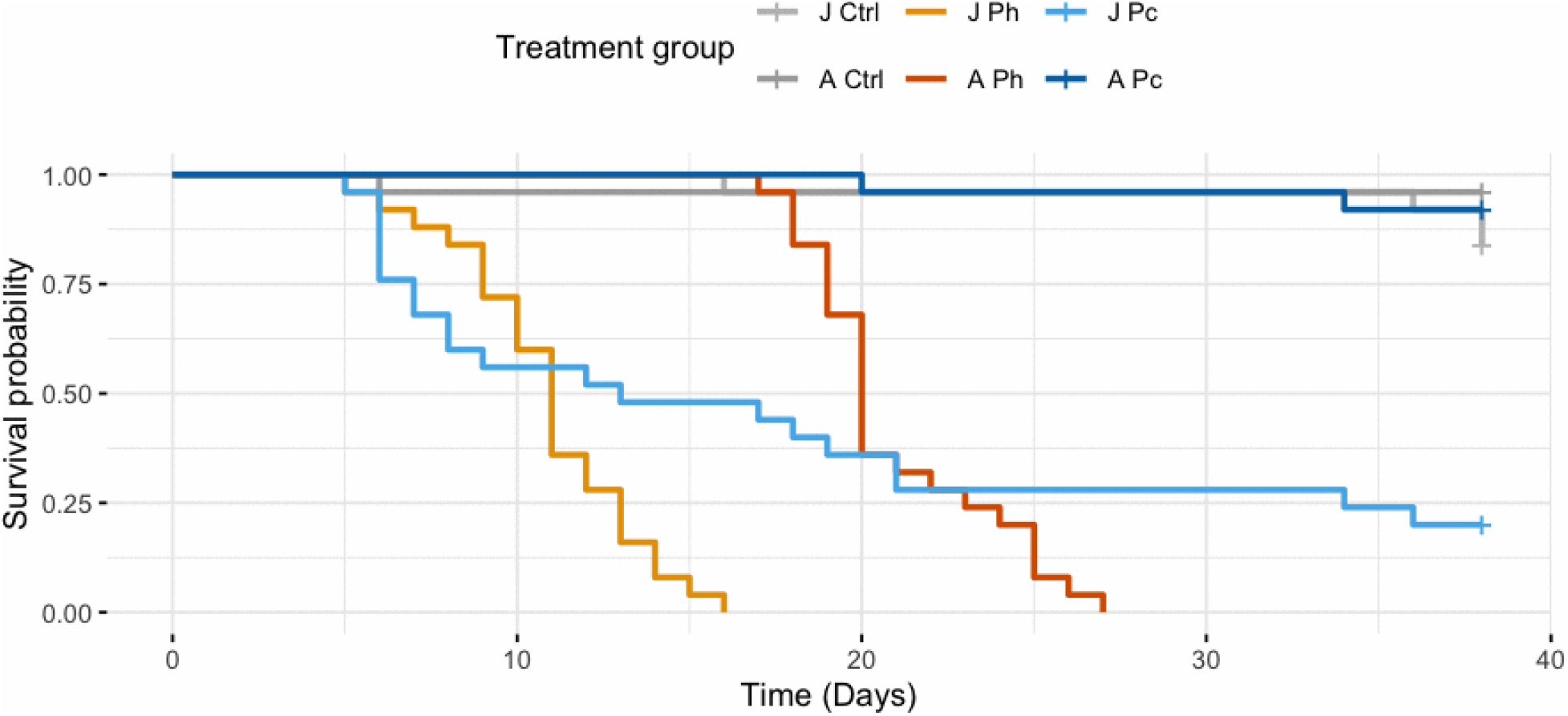
Kaplan-Meier survival probability (y-axis) as a function of number of exposure days (x-axis). Results for each treatment group are shown (Juvenile control, light grey; Adult control, dark grey; Juvenile *P. hermaphrodita*, light orange; Adult *P. hermaphrodita*, dark orange; Juvenile *P. californica*, light blue; Adult *P. californica*, dark blue).

## Discussion

It was observed that both *P. hermaphrodita* and *P. californica*, two species commercialized as biocontrol products targeting pest slugs (such as *D. reticulatum*), increase mortality in a Pacific coast-native and non-target species of slug, *A. columbianus*, in laboratory conditions (Fig 5). Survival probability was shown to differ significantly based on the life stage of the host organism (juvenile or adult stage *A. columbianus*) (Table 1). These findings suggest that *Pellioditis* spp. nematodes could have negative effects on native terrestrial gastropod biodiversity and should be used with extreme caution to reduce potential non-target species encounters.

The two *Pellioditis* species used in this infectivity trial, *P. hermaphrodita* (strain DL309) and *P. californica* (strain DL320), differed significantly in their effect on *A. columbianus* survival in a laboratory setting (Table 1). All slugs that were exposed to *P. hermaphrodita* experienced complete mortality by the end of the trial, regardless of juvenile or adult life stage. In contrast, while *P. californica* caused substantial mortality in the juvenile group, this treatment never led to complete mortality in either life stage, with adult slugs experiencing very little death comparable to that of the control treatments (Fig 4, 5). Pairwise log-rank comparisons found a significant difference in survival probability between adult *P. hermaphrodita* and *P. californica* treatments, but not between the corresponding juvenile groups (Table 1). The differential survival of *A. columbianus* observed here is consistent with previous findings comparing the infectivity of four *Pellioditis* species in pest slug *D. reticulatum*, in which *P. hermaphrodita* caused mortality at a faster rate as compared to *P. californica* [11].

Results of this study demonstrated significant differences in survival probability between life stages (Table 1). In both *Pellioditis* spp. treatments, the *A. columbianus* juvenile stage experienced mortality at a faster rate than the corresponding adult groups (Fig 4, 5), suggesting that smaller or less developed individuals are more likely to experience mortality when exposed to slug-killing nematodes. These results are consistent with size and life stage-dependent infectivity patterns observed across other species, including *Arion ater* and *Arion lusitanicus* [17, 24]. This has important implications for how local gastropod communities may be impacted by biocontrol nematodes, where differential survival depending on life stage could alter population dynamics. This builds on a growing body of work providing strong rationale for comprehensive non-target testing of multiple life stages to thoroughly understand potential impacts.

This study provides evidence of a potential non-target effect of *Pellioditis* biocontrol nematodes on the Pacific-coast native *A. columbianus*. This expands on previous infectivity testing of non-targets in this lab, which observed *P. hermaphrodita* to cause lethality in the native *M. fidelis* in laboratory infectivity trials [14]. While *P. hermaphrodita* infectivity has been studied with regards to use in Europe and determined not to pose a significant risk to non-targets [9], *P. californica* has less information available. In both species, there is little knowledge of potential non-target effects in the U.S. specifically. Additionally, in contrast to European studies where common hedgerow gastropods are identified [13], it is not clear which species in North American ecosystems will be most at risk of exposure. Future work is needed to understand the context in which slug-killing nematodes might be used in the U.S., and which native terrestrial gastropods would be most likely to encounter these applications. The findings of this laboratory infectivity trial suggest that commercial nematode products, such as Nemaslug®, may have significant non-target impacts in native gastropod communities of the American Pacific Northwest. However, further studies are needed to understand if this result is replicable in a more natural setting, such as a mesocosm, to account for environmental variables that are not present in the lab assay. Furthermore, while this study uses a mortality-based assay, there could be other non-lethal effects on non-target organisms, such as reduced growth or reduced immune system function, that should be explored to fully understand the ecological impact of using these nematodes as biocontrol.

In conclusion, this infectivity trial demonstrated not only the susceptibility of *A. columbianus* to infection and mortality by *P. hermaphrodita* and *P. californica*, but also that susceptibility may depend on life stage of the host organism. Current regulatory frameworks in the U.S. may not adequately capture potential impact associated with life stage or specific regional non-target species. We therefore urge extreme caution in implementing biological control strategies utilizing gastropod-killing nematodes, and advocate for comprehensive non-target testing of many species and life stages to better understand how they may impact local gastropod biodiversity.

## Acknowledgements

The authors express gratitude to Brenna Prevelige and Jazlee Crowley for assisting with field collection. We’d also like to thank Yanming Di for advice on statistical analysis.

## References

1. Lefcheck JS, Byrnes JEK, Isbell F, Gamfeldt L, Griffin JN, Eisenhauer N, et al. Biodiversity enhances ecosystem multifunctionality across trophic levels and habitats. Nat Commun. 2015 Apr 24;6(1):6936.

2. Salisbury S. The cost of slugs to the grass seed industry in the Willamette Valley. [presentation]. Salem (OR): Oregon State Legislature; 2015 May.

3. Lydeard C, Cowie RH, Ponder WF, Bogan AE, Bouchet P, Clark SA, et al. The global decline of nonmarine mollusks. BioScience. 2004 Apr 1;54(4):321–30.

4. Meyer WM, Ostertag R, Cowie RH. Influence of terrestrial molluscs on litter decomposition and nutrient release in a Hawaiian rain forest. Biotropica. 2013 Nov;45(6):719–27.

5. IUCN Red List of Threatened Species [Internet]. Gland (Switzerland): International Union for Conservation of Nature; [cited 2026 Jan 22]. Available from: https://www.iucnredlist.org

6. Christensen CC, Cowie RH, Yeung NW, Hayes KA. Biological control of pest nonmarine molluscs: A Pacific perspective on risks to non-target organisms. Insects. 2021 Jul;12(7):583.

7. Régnier C, Fontaine B, Cowie RH, Bouchet P. Measuring the Sixth Extinction: what do mollusks tell us? The Nautilus [Internet]. 2017 [cited 2024 Jul 28]. Available from: https://hal.science/hal-03864260

8. Gerlach J, Barker GM, Bick CS, Bouchet P, Brodie G, Christensen CC, et al. Negative impacts of invasive predators used as biological control agents against the pest snail *Lissachatina fulica*: the snail *Euglandina* ‘*rosea*’ and the flatworm *Platydemus manokwari*. Biol Invasions. 2021 Apr 1;23(4):997–1031.

9. Rae R, Sheehy L, McDonald-Howard K. Thirty years of slug control using the parasitic nematode *Phasmarhabditis hermaphrodita* and beyond. Pest Management Science. 2023;79(10):3408–24.

10. Stevens G, Lewis E. Status of entomopathogenic nematodes in integrated pest management strategies in the USA. In: Hajek AE, editor. Biocontrol agents: entomopathogenic and slug parasitic nematodes [Internet]. Wallingford (UK): CABI; 2017 [cited 2026 Jan 22]. p. 289–311. Available from: 10.1079/9781786390004.0289

11. Mc Donnell RJ, Colton AJ, Howe DK, Denver DR. Lethality of four species of *Phasmarhabditis* (Nematoda: Rhabditidae) to the invasive slug, *Deroceras reticulatum* (Gastropoda: Agriolimacidae) in laboratory infectivity trials. Biological Control. 2020 Nov 1;150:1043–49.

12. Castle GD, Mills GA, Gravell A, Jones L, Townsend I, Cameron DG, et al. Review of the molluscicide metaldehyde in the environment. Environmental Science: Water Research & Technology. 2017;3(3):415–28.

13. Wilson MJ, Hughes LA, Hamacher GM, Glen DM. Effects of *Phasmarhabditis hermaphrodita* on non-target molluscs. Pest Management Science. 2000;56(8):711–6.

14. Denver D, Howe DK, Colton AJ, Richart CH, Donnell RJM. The biocontrol nematode *Phasmarhabditis hermaphrodita* infects and increases mortality of *Monadenia fidelis*, a non-target terrestrial gastropod species endemic to the Pacific Northwest of North America, in laboratory conditions. PLOS ONE. 2024 Mar 21;19(3):e0298165.

15. Harper AB. The banana slug: A close look at a giant forest slug of Western North America. Aptos, CA: Bay Leaves Press; 1988.

16. Wilson MJ, Glen DM, George SK. The rhabditid nematode *Phasmarhabditis hermaphrodita* as a potential biological control agent for slugs. Biocontrol Science and Technology. 1993 Jan 1;3(4):503–11.

17. Speiser B, Zaller JG, Neudecker A. Size-specific susceptibility of the pest slugs *Deroceras reticulatum* and *Arion lusitanicus* to the nematode biocontrol agent *Phasmarhabditis hermaphrodita*. BioControl. 2001 Sep 1;46(3):311–20.

18. Grewal SK, Grewal PS, Hammond RB. Susceptibility of North American native and nonnative slugs (Mollusca: Gastropoda) to *Phasmarhabditis hermaphrodita* (Nematoda: Rhabditidae). Biocontrol Science and Technology. 2003 Feb 1;13(1):119–25.

19. Rae R, Verdun C, Grewal PS, Robertson JF, Wilson MJ. Biological control of terrestrial molluscs using *Phasmarhabditis hermaphrodita*—progress and prospects. Pest Management Science. 2007;63(12):1153–64.

20. Grannell A, Cutler J, Rae R. Size-susceptibility of *Cornu aspersum* exposed to the malacopathogenic nematodes *Phasmarhabditis hermaphrodita* and *P. californica*. Biocontrol Science and Technology. 2021 Nov 2;31(11):1149–60.

21. Schurkman J, Ley ITD, Dillman AR. Dose dependence of *Phasmarhabditis* isolates (*P. hermaphrodita, P. californica, P. papillosa*) on the mortality of adult invasive white garden snails (*Theba pisana*). PLOS ONE. 2022 Jul 22;17(7):e0270185.

22. Schurkman J, Dodge C, Donnell RM, Ley ITD, Dillman AR. Lethality of *Phasmarhabditis* spp. (*P. hermaphrodita, P. californica*, and *P. papillosa*) nematodes to the grey field slug *Deroceras reticulatum* on Canna lilies in a lath house. Agronomy [Internet]. 2021 Dec 22 [cited 2026 Jan 24];12(1):20. Available from: https://www.mdpi.com/2073-4395/12/1/20

23. Schurkman J, Ley ITD, Dillman AR. Lethality of three *Phasmarhabditis* spp. (*P. hermaphrodita, P. californica*, and *P. papillosa*) to *Succinea* snails. Agriculture [Internet]. 2022 Jun 9 [cited 2026 Jan 24];12(6):837. Available from: https://www.mdpi.com/2077-0472/12/6/837

24. Grimm B. Effect of the nematode *Phasmarhabditis hermaphrodita* on young stages of the pest slug *Arion lusitanicus*. J Molluscan Stud. 2002 Feb 1;68(1):25–8.

25. Pearson AK, Pearson OP, Ralph PL. Growth and activity patterns in a backyard population of the banana slug, *Ariolimax columbianus*. Veliger. 2006;48(3):143–50.

26. Ley ITD, Kiontke K, Bert W, Sudhaus W, Fitch DHA. *Pellioditis pelhamensis* n. sp. (Nematoda: Rhabditidae) and *Pellioditis pellio* (Schneider, 1866), earthworm associates from different subclades within *Pellioditis* (syn. *Phasmarhabditis* Andrássy, 1976). PLOS ONE. 2023 Sep 6;18(9):e0288196.

27. Orozco RA, Lee MM, Stock SP. Soil sampling and isolation of entomopathogenic nematodes (Steinernematidae, Heterorhabditidae). J Vis Exp. 2014 Jul 11;(89):52083.

